# Honeybee genetics shape the strain-level structure of gut microbiota in social transmission

**DOI:** 10.1101/2020.12.17.423353

**Authors:** Jiaqiang Wu, Haoyu Lang, Xiaohuan Mu, Zijing Zhang, Qinzhi Su, Xiaosong Hu, Hao Zheng

**Author notes:** These authors contributed equally (Jiaqiang Wu, Haoyu Lang).

## Abstract

Honeybee gut microbiota transmitted via social interactions are beneficial to the host health. Although the microbial community is relatively stable, individual variations and high strain-level diversity have been detected across honeybees. Although the bee gut microbiota structure is influenced by environmental factors, the heritability of the gut members and the contribution of the host genetics remains elusive. Considering bees within a colony are not readily genetically identical due to the polyandry of queen, we hypothesize that the microbiota structure can be shaped by host genetics. We used shotgun metagenomics to simultaneously profile the microbiota and host genotypes of individuals from hives of four different subspecies. Gut composition is more distant between genetically different bees at both phylotype- and “sequence-discrete population”-level. We then performed a successive passaging experiment within colonies of hybrid bees generated by artificial insemination, which revealed that the microbial composition dramatically shifts across batches of bees during the social transmission. Specifically, different strains from the phylotype of *Snodgrassella alvi* are preferentially selected by genetically varied hosts, and strains from different hosts show a remarkably biased distribution of single-nucleotide polymorphism in the Type IV pili loci. A genome-wide association analysis identified that the relative abundance of a cluster of *Bifidobacterium* strains is associated with the host glutamate receptor gene that is specifically expressed in the bee brain. Finally, mono-colonization of *Bifidobacterium* with a specific polysaccharide utilization locus impacts the expression and alternative splicing of the *gluR-B* gene, which is associated with an altered circulating metabolomic profile. Our results indicated that host genetics influence the bee gut composition, and suggest a gut-brain connection implicated in the gut bacterial strain preference. Honeybees have been used extensively as a model organism for social behaviors, genetics, and gut microbiome. Further identification of host genetic function as shaping force of microbial structure will advance our understanding of the host-microbe interactions.

## Background

It is becoming increasingly clear that most animals are universally inhabited by microbial communities in their guts. These host-associated microbiomes provide considerable benefits to the host through different functions, and shape the host’s ecology and evolution[1] Microbiomes in animals are often acquired at birth, as well as through contact with others and the environment. Considering the importance of the microbiome, it is thus crucial to understand mechanisms that select, retain, and transfer the commensal microbes and the impact of specific constituents on host biology. Numerous environmental factors, including lifestyle, diet, disease, geography and medications have an influence on the gut microbiota[2–4]. In addition to the external factors, host genetics has recently been proposed as a determinant of gut microbial composition[5]. Studies searched for associations between host alleles and gut microbiota structure of human, mice and other animal models[6–8]. Different genetic loci have emerged as influential in accounting for interindividual variation in microbiomes, specifically, genes with roles in host metabolism and immune systems implicated in disease have been studied for their impact on the microbiome[9–12]. In addition, a suite of taxa are considered more heritable than others, suggesting that host genetic variation can explain the levels of certain gut members. Particular taxa, such as Christensenellaceae and methanogens, are linked to the host lean phenotype, suggesting the important functions and underlying host-microbe interactions of these highly heritable gut members[13–15]. Host gene-microbiome cross-talk has been surveyed for humans, as well as other animals, yet it is challenging for the association of specific host alleles with microbiome traits. This is partly due to the gut community structure being strongly and rapidly affected by a variety of environmental factors, making comparisons of animals in controlled laboratory settings to minimize environmental influences difficult, limiting our ability to extrapolate host-microbe interactions. Moreover, the genome-wide association study (GWAS) for hosts with a relatively large genome size with complex microbiota dominated by thousands of bacterial species is costly[16]. Therefore, experimental systems with simple microbiota composition that are able to be controlled in designated conditions are required, in order to obtain a better understanding of the processes that determines microbiome structure and transmission dynamics.

Honeybees possess a simple and host-restricted gut microbiota that is commonly detected in bees that are even derived from different locations[17, 18]. This community contains only 5–8 core bacterial phylotypes (with >97% sequence similarity in the 16S rRNA gene), whereas multiple “sequence-discrete populations” (SDPs; with ~90% genomic average nucleotide identity) and strains have been defined in each phylotype, indicating a strain-level variation in the bee gut[19, 20]. In particular, although only one phylotype has been identified for the core gut member *Snodgrassella*, a high strain-level diversity and host-specific pattern has been indicated[21, 22]. Multiple strains from all core members of bee gut bacteria have been isolated, and the ease of rearing microbiota-free (MF) bees facilitates a functional screen of individual gut members using gnotobiotic bees inoculated with defined communities[23]. Thus, the bee gut community provides an excellent model for studies on the processes that govern the assembly of microbiota. To date, studies have focused on the roles of bee gut microbiota, such as their effects on host nutrition, weight gain, endocrine signaling and immune functions[24, 25]. However, the knowledge of the effectors that affect the composition and transmission of the honeybee gut microbiota is quite limiting.

It has been documented that the characteristic bee gut microbiota is mainly acquired from the other nest mates through social contact, and the bee gut microbiota are vertically inherited inside the colony[26]. Although the western honeybee (*Apis mellifera*) gut microbiota are relatively stable across colonies from different geographic locations, individuals with varying host behaviors and physiologies, such as age and caste, possess distinctive gut compositions[27, 28]. Moreover, gut compositions vary in individuals even from the same colony, with a high diversity in strain composition for the core gut members being observed in particular[21]. Worker bees from the same hive are not readily genetically identical, since queen bees are polyandrous and mate with an average of 12 males to produce daughters of mixed paternity[29, 30]. Indeed, a previous study showed that colonies with genetically diverse populations exhibit a higher microbiota diversity, strongly suggesting a host genetic impact on gut community[31]. We hypothesized that host genetics is a shaping force of the bee gut microbial diversity, especially the stain-level variation and the dynamics of microbiota transmission along generations.

Here, using shotgun metagenomics, we simultaneously profiled the strain-level bacterial community and host genotype for individual pure race of bees and hybrids generated through artificial insemination. It was observed that the abundances of most core gut members were influenced by host genetics. A longitudinal study of the structure and dynamics of the gut communities found that specific taxa were selected by the host during the successive passage across genetically varied generations. Heritability analysis and a GWAS on gut composition identified a significant association between *Bifidobacterium* and the glutamate receptor gene preferably expressed in the bee brain. Furthermore, the brains of bees colonized with the heritable *Bifidobacterium* displayed a differential expression level and differential splicing events of the *gluR-B* gene, which may be partly explained by a glycan utilization gene cluster specific to the highly heritable strains.

## Methods

### Honeybee management and samples collection

All honey bees (*Apis mellifera*) were bred and maintained in the apiary of Jilin Province Institute of Apicultural Science. The purebred swarms were established since 1980s when the queens were imported, and all breeds were conserved by artificial insemination each year for long-term genetic studies. Four different subspecies, namely OH, AF, YF, and SK were used in this study. We set up one hive for OH and AF each, and two hives of YF and SK each, and both the purebred and hybrid colonies were constructed in July, 2018. For the hybrid colonies, virgin queens of OH and drones of AF or YF were mated by artificial insemination to produce hybrids O-A and O-Y, respectively. The queens were singly mated by instrumentally inseminated with semen harvested from a single drone. Then the inseminated queens were placed into nucleus colonies with ~300 founding workers, named O-A’ and O-Y’ here. When the queens started laying eggs, each colony was monitored daily. To control the age of sampled bees, one-day-old workers were obtained by moving frames containing late-stage pupae from colonies into an incubator (35 °C, 50% relative humidity) overnight. The newly emerged adults in the incubator were individually marked on their thoraces with a spot of color paint and were then placed back into their original hives, and all individuals were collected when they are 15 day old. When the first batch of bees (B1) were collected, the second batch (B2) was set up as described above. In total, three batches of bees (B1–B3) were labeled and sampled consecutively (Fig. 2a). Notably, before the emergence of B3, it had been more than 50 days since the founding workers introduced, thus, all O-A’ and O-Y’ bees had died.

### DNA extraction and shotgun sequencing

All bee individuals of either purebred or hybrid were sampled exactly 15 day after the emergence. Total genomic DNA of both the bee host and gut microbiota was extracted from the whole gut homogenate using CTAB method as previously described[24]. In brief, the whole gut was resuspended in 728 μl CTAB buffer and 20 ul 20mg/ml proteinase K (TransGen Biotech) in a 2-ml tube containing 500 μl of sterile zirconia beads Ø 0.1 mm (BioSpec, Bartlesville, OK, USA). Cells were lysed by running MO BIO Vortex Genie for 3 min, and were centrifuged at 12,000×g for 5 min. The supernatant was transferred to a clean 1.5-ml tube and incubated at 56°C for 30 min. After cooling at room temperature for 3 min, 5 μl RNase A (TransGen Biotech) was added, and incubated at 37°C for 30min. Samples were mixed with 400 μl Phenol:Chloroform:Isoamyl alcohol, and centrifuged at 12,000×g for 5 min. The supernatants were transferred to new 1.5-ml tubes and 50 μl sodium acetate (pH 5.5) and 500 μl isopropanol was added. After centrifuging at 17,000×g for 30 min, the pellets were washed twice with 70% ethanol. The DNA pellets were then dissolved in TE buffer (pH 8.0). DNA samples were sent to Guangdong Magigene Biotechnology Co.,Ltd. (Guangzhou, China) for shotgun metagenome sequencing. Sequencing libraries were generated using NEBNext UltraTM II DNA Library Prep Kit for Illumina (New England Biolabs, MA, USA), and the library quality was assessed on Qubit 3.0 Fluorometer (Life Technologies, Grand Island, NY) and Agilent 4200 (Agilent, Santa Clara, CA) system. The libraries were then sequenced on the Illumina HiSeq X-ten platform with 150 bp paired-end reads.

### Mapping and Variant Calling on Honeybees

The raw data obtained from the Illumina HiSeq sequencing platform was preprocessed by Readfq: reads with low quality bases above 40 bp (quality value ≤ 38), with N bases >10 bp were removed, and sharing the overlap >15 bp with the adapters were all removed. In addition, reads were quality controlled with Fastp using default parameters to generate clean data for downstream analysis. ~10Gb of sequences per sample were obtained for subsequent analyses. The quality-controlled reads were mapped to the honeybee reference genome (version Amel_HAv3.1; GenBank assembly accession: GCA_003254395.2) using BWA-MEM algorithm with the option “-t 4, −R, −M”. We then sorted the alignments according to mapping coordinates and marked PCR duplicates using Picard Tools, and the duplications were removed using the SAMtools program to create one BAM file for each bee sample. To improve mapping quality, we set minimum mapping quality = 1. We defined depth of coverage as the number of mapped bases divided by the total length of the reference genome, and the coverage of more than 90% of the samples are above 20X. As a basic quality control metric, 95%-100% (median 99.14%) of mapped reads per sample were properly in pairs.

We called variants using the Genome Analysis Toolkit v4.0.3.0 following germline short variant discovery pipeline of the best-practices. In brief, HaplotypeCaller module with default parameters was used to calculate variant calls independently for each BAM file and generate one gVCF file per sample. Then, gVCFs were merged into GenimicsDB across all samples by GenomicsDBImport module for the following joint genotyping, which was performed using GenotypeGVCF on all samples to create a single VCF file for the whole population. To improve the variants quality, we filtered out false positive SNPs with VCFtools and checked the dataset by VariantQC. Finally, only sites with a minor allele frequency between 0.05 and 0.95 were kept. The quality of the final set of SNPs was assessed by calculating the ratios of transition to transversion (Ti/Tv ratio), a diagnostic parameter to examine the quality of SNP identification. The ratios distributed between 4.17 and 4.29 in our population, which indicates no excess of false-positive SNPs.

### Honeybee population genetics analyses

To investigate genetic relatedness of purebred bees including OH, YF, AF and SK, the gVCF file of *Apis cerana* (version ACSNU-2.0) was merged with the population VCF dataset of *A. mellifera* created above. We then measured the raw genetic distance using the ‘SNPRelate’ package based on the number of shared alleles between each individual sample. The distance matrix was used to estimate a tree using neighbor-joining method implemented in R package ‘ape’ and the tree was visualized using iTol. PLINK was used to generate a clear population VCF dataset with only biallelic loci, all variants with missing call rates exceeding 0.05 were filtered out. Markers with a Hardy-Weinberg Equilibrium p-value < 0.0001 and individuals with less than 0.05 missing genotype data were excluded. In total, we identified 2,255,909 high quality variants (average value = 20,482 per site) across the 57 purebred bees. The numbers of SNVs from all samples presented across *A. mellifera* chromosome 1 to 16 in 100 kb consecutive windows are shown in Fig.S2. We also processed the same pipeline to acquire variant files of hybrid and founding worker bees (Fig. 2a), which resulted in 2,444,291 variants (average value = 31,690 per site) across the 68 individuals. We then ran ADMIXTURE on the resulting SNPs to estimate the genetic ancestry of each sample. The unsupervised analysis was performed with the number of hypothetical ancestral components (*K*) ranging from 2 to 5, and the five-fold cross-validation (CV) error values were estimated in ADMIXTURE at each *K* value.

### Isolation and genome sequencing of gut bacteria

Bacterial strains were isolated from the guts of *A. mellifera* collected in Jilin, China during July 2018 (Additional file 1: Dataset S1). The dissected guts were directly crushed in 20% (vol/vol) glycerol and frozen at −80 °C after sampling. The glycerol stocks were plated on heart infusion agar supplemented with 5% (vol/vol) defibrinated sheep’s blood (Solarbio, Beijing, China), MRS agar (Solarbio, Beijing, China) or TPY agar (Solarbio, Beijing, China) incubated at 35 °C under a CO_2_-enriched atmosphere (5%). Genomic DNA of bacterial isolates was extracted using the CTAB buffer method as previously described [32]. Total genomic DNA of the isolate was sequenced on the Illumina Nova6000 platform from the 150 bp paired-end libraries, and the quality-controlled reads were assembled with the SOAPdenovo2 genome assembler. The completeness of genomes was assessed by CheckM. Whole-genome average nucleotide identity (ANI) was calculated using FastANI. The genomes were annotated with the Prokka software and phylogenetic trees were reconstructed based on the whole-genome sequence information by PhyloPhlAn. All genome assemblies obtained in this study were deposited at DDBJ/EMBL/GenBank, and the accession numbers are listed in Additional file 1: Dataset S1.

### Metagenomic analysis

The SDP- and strain-level community structure of each sample was profiled following the Metagenomic Intra-Species Diversity Analysis System (MIDAS) pipeline. Firstly, a custom genomic database for taxonomy classification and strain SNP calling was built using genomes of 116 new bee gut bacterial isolates obtained in this study together with 289 published genomes isolated from *Apis* and *Bombus* for the downstream MIDAS analysis (Additional file 1: Dataset S1). According to the MIDAS pipeline instruction, genomes with an exact 95% ANI pairwise similarity were grouped into one species cluster, which were defined as “MIDAS-taxonomy” here. Representative genomes from each species cluster were chosen to maximize its average nucleotide identity at the 30 universal genes to other members of the species cluster, and they were used for detecting core-genome SNPs. Next, a database of 15 universal single-copy gene families was built to estimate the abundance of the species clusters from the shotgun metagenomes. Gene families were selected by MIDAS based on their ability to accurately recruit metagenomic reads as well as being universal and single copy.

Ellegaard et al. identified that multiple sequence-discrete populations (SDPs) are existing within the core phylotypes of the gut community of *A. mellifera*. Our analyses of the isolate genome phylogeny and genomic pairwise ANI confirmed the existence of most of the SPDs (Figure S1). However, we noticed that the pairwise ANI within the predefined SDP “Bifido-1” are much higher than the expected threshold (95%). Moreover, our previous work has documented that strains from “Bifido-1” show dramatically various capacity in polysaccharide digestion[32], indicating that they might belong to different bacterial populations. Here, we included 28 new bifidobacteria isolates in the analyses of genome phylogeny and pairwise genomic ANI. We identified that “Bifido-1” can be further divided into four SDPs, which form discrete clades and show similar metabolic profiles [32]. Thus, the bacterial genomes were integrated into 76 MIDAS-species clusters and they were further grouped into 17 SDPs for the five core phylotypes and *Bartonella apis*. There are also 20 SDPs for the other non-core members included in the database for the taxonomy classification. The taxonomic annotations of the MIDAS custom database are shown in Additional file 1: Dataset S1.

Before the taxonomic profiling, we removed host-derived reads from metagenomic clean data with KneadData, which maps the reads to the honeybee reference genome (Amel_HAv3.1) with default parameters and performs quality control. To compute relative abundance of SDPs, coverage and prevalence from metagenomic sequencing, we ran ‘species’ module of the ‘run_midas.py’ script and ‘merge_midas.py’ script with our custom bacterial genome database. MIDAS aligned reads to the database of universal single-copy marker gene sequences for each of the species cluster within our custom database using HS-BLASTN to estimate the abundance of species clusters for each sample. Local alignments that cover < 70% of the read or fail to satisfy the gene-specific species-level percent identity cut-offs were discarded, and we assigned each uniquely mapped read to a species cluster according to its best hit. Then, the coverage and relative abundance of each species cluster across samples were estimated based on the number of mapped reads.

To have insight into community diversity of samples of inter- and intra-subspecies of honeybees, we compared both phylotype- and SDP-level composition profiles by measuring Bray-Curtis distance using the ‘vegan’ package. We used Wilcoxon tests to evaluate the dissimilarity for each subject. Then, we compared the relative abundance of all phylotypes and SDPs for different subspecies of bees, and we used Kruskal-Wallis test to determine whether it is statistically significant different between honeybee groups.

### *Snodgrassella alvi* phylotype-specific strain analysis

We used the ‘single-nucleotide-polymorphism prediction’ function of the MIDAS pipeline to profile SNPs of *Snodgrassella* in metagenomic data for each individual bee against the representative genome. In brief, we ran the ‘snps’ module of the ‘run_midas.py’ script to map metagenomic reads to the reference genome of *Snodgrassella alvi*. Strain M0112 was selected as it meets the coverage requirement and possesses the greatest nucleotide identity to other members of the species. Metagenomic reads were aligned to the reference genomes using Bowtie2, with default MIDAS mapping thresholds: global alignment, mapping percent identity ≥ 94.0%, sequence quality ≥ 20, alignment coverage ≥ 0.75, and mapping quality ≥ Pileups of each sample were generated using SAMtools and the nucleotide variation statistics were then counted at each genomic site. After running this script for each sample, we pooled all samples and performed core-genome SNP calling using the ‘snps’ module of ‘merge_midas.py’ script, and the core-genome SNP matrices for *S. alvi* were produced for the comparative analyses of nucleotide variation across genomic sites and metagenomic samples of different bee groups. Specifically, bi-alleles genomic sites present in more than 95% of strains and with a minimum prevalence frequency of 1% were identified as core SNPs using the ‘--core-snps’ parameter. Based on the raw alignments, MIDAS reported the read coverage for each site in the reference genome for each metagenomic sample. The frequencies of both silent and missense allele per genomic site per sample in protein-coding genes were used to indicate the strain-level variation among different individuals as previously described [33, 34].

### 16S rRNA high throughput sequencing

The V4 region of 16S rRNA gene was amplified (primers 515F and 806R) with barcodes. All PCR reactions were carried out with 15 μL of Phusion® High-Fidelity PCR Master Mix (New England Biolabs); 0.2 μM each of forward and reverse primers, and 10 ng template DNA. PCR products was purified with Qiagen Gel Extraction Kit (Qiagen, Germany). Sequencing libraries were generated using NEBNext® UltraTM II DNA Library Prep Kit for Illumina^®^ (New England Biolabs, MA, USA) and index codes were added. The library quality was assessed on the Qubit 2.0 Fluorometer (Thermo Fisher Scientific, MA, USA). The library was sequenced on an Illumina Nova6000 platform with 250 bp paired-end reads, and at least 30,000 sequences were obtained for each individual sample. Fastp [35] was used to control the quality of the raw data by sliding window (window size=4 bp, mean quality=20. The primers were removed using Cutadapt software[36] according to the primer information at the beginning and end of sequences to obtain clean reads. Mate-pair merging, OTU picking, chimera elimination and taxonomy classification were performed using Mothur version 1.40.5 following the MiSeq standard operating procedure[37]. 16S rRNA gene sequences were clustered into Operational Taxonomic Units (OTUs) at a similarity cutoff value of 99%. We used a curated database for bee gut microbiota based on SILVA database for the classification[38]. The subsequent analysis was based on normalized OTU table using the ‘phyloseq’ package[39], and Bray-Curtis diversity was calculated with the ‘vegan’ package[40].

### Heritability calculations

The heritability of each gut bacteria taxa is defined as the proportion of total variance due to genetic effects as previously described[11].The heritability was calculated using additive effect model in HIBLUP V1.3.1 under Genomic Best Linear Unbiased Prediction framework[41]. The VCF datasets of individuals from OH, YF, AF, SK, O-A and O-Y groups were used here. Variants in the host genome were quality controlled, sites with a minor allele frequency < 0.05 or > 0.95 or failed in the Hardy-Weinberg test at 0.0001 were removed, which results in 2,861,994 informative SNPs across 102 bee samples and the average quality value is 49,095. A combination of HE Regression algorithm and Average Information algorithm was used to obtain an efficient and robust variance component estimation. Grouping of hives and kinship among individuals are used as random effects.

### Bacterial Associations to Host Genetic Variation

We performed GWAS analysis for the relative abundances of each core bacteria phylotype and SDP using rMVP v1.0.0 with correcting for population structure. We used the SNP profiles of 102 individuals from OH, YF, AF, SK, O-A and O-Y group of bees from the clear VCF dataset. All sites were filtered with PLINK software as described above. General Linear Model (GLM) and Mixed Linear Model (MLM) were both used for the host SNP-microbe association tests, and we used ‘GEMMA’ method to analyze the variance components. The relatedness matrix, measured as the genetic similarity between individual bees, was used to estimate random effects. For all samples, SNPs and the top three PCs were used as fixed effects in MLM, and the top five PCs were used in GLM. The p-values were firstly set at 0.05 for each association test, and then the Benjamini-Hochberg corrected p-value threshold for all SNPs was used to control for false-positive error rates deriving from multiple testing at the genome-wide level.

### Targeted metabolomics for GABA in bee brains

Brain tissues dissected from MF and ‘Bifido1.4’-colonized bees were sent to Biotree Biotech Co. Ltd. (Shanghai, China) for targeted metabolomics analysis of GABA. Six brain tissues from one treatment group were put into one tube and centrifuged (2400 g × 1 min at 4 °C). 100 μL acetonitrile containing 0.1% formic acid and 20 μL ultrapure water were added and the tubes were vortexed thoroughly. Tissue cells were sonicated in ice-water bath for 30 min, followed by subsiding at −20 °C for 2 h. Supernatants were collected after centrifugation (14,000 g × 10 min at 4 °C). 20 μL of supernatant were transferred to a new vial followed by incubation for 30 min after the addition of 10 μL sodium carbonate solution (100 mM) and 10 μL 2% benzoyl chloride acetonitrile. Then 1.6 μL internal standard and 20 μL 0.1% formic acid were added, and the samples were centrifuged (14,000 g × 5 min at 4 °C). 40 μL of the supernatants were transferred to an auto-sampler vial for downstream UHPLC-MS/MS analysis. 4-aminobutyric acid (Sigma-Aldrich) was used for the construction of calibration standard curve.

The UHPLC separation was carried out using an ExionLC System (AB SCIEX; MA, USA), and the samples was analyzed on the QTRAP 6500 LC-MS/MS system (AB Sciex; Framingham, MA, USA). 2 μL of samples was directly injected onto an ACQUITY UPLC HSS T3 column (100 × 2.1 mm × 1.8 μm; Waters; Milford, Ma, USA). The column temperature was set at 40 °C, and the auto-sampler temperature was set at 4 °C. Chromatographic separation was achieved using a 0.30 ml/min flow rate and a linear gradient of 0 to 2% B within 2 min; 2%–98% B in 9 min, followed by 98% B for 2 min and equilibration for 2 min. Solvent A is 0.1% formic acid and solvent B is acetonitrile. For all multiple reaction monitoring (MRM) experiments, 6500 QTrap acquisition parameters were as follows: 5000 V Ion-spray voltage, curtain gas setting of 35 and nebulizer gas setting of 60, temperature at 400 °C. Raw data were analyzed using Skyline[42].

### Quasi-targeted metabolomics of hemolymph

Honeybee hemolymph was collected using a 10 μL pipettor from the incision above the median ocellus of bees as previously described[24]. A minimum of 50 μL of hemolymph was collected from10 bees into a 1.5-mL centrifuge tube. During the collection process, tubes are preserved on dry ice and subsequently stored at −80 °C until instrumental analysis. Hymolymph metabolites were determined by quasi-targeted metabolomics using HPLC-MS/MS. 50 μL of hemolymph samples were mixed with 400 μL prechilled methanol by vortexing. All samples were incubated on ice for 5 min and then centrifuged at 15,000 × g, at 4°C for 10 min. The supernatant was diluted to a final concentration containing 53% methanol by LC-MS grade water. The samples were then transferred to a fresh vial and centrifuged at 15,000 × g, 4°C for 20 min. Finally, the supernatant was injected into the LC-MS/MS system, and the analyses were performed using an ExionLC AD system (SCIEX) coupled with a QTRAP 6500+ mass spectrometer (SCIEX). Samples were injected onto a BEH C8 Column (100 mm × 2.1 mm × 1.9 μm) using a 30-min linear gradient at a flow rate of 0.35 mL/min for the positive polarity mode. Eluent A was 0.1% formic acid-water and eluent B is 0.1% formic acid-acetonitrile. The solvent gradient was set as follows: 5% B, 1 min; 5-100% B, 24.0 min; 100% B, 28.0 min;100-5% B, 28.1 min;5% B, 30 min. QTRAP 6500+ mass spectrometer was operated in positive polarity mode with curtain gas of 35 psi, collision gas of Medium, ion spray voltage of 5500V, temperature of 500°C, ion source gas of 1:55, and ion source gas of 2:55. For negative ion mode, samples were injected onto aHSS T3 Column (100 mm × 2.1 mm) using a 25-min linear gradient at a flow rate of 0.35 mL/min. The solvent gradient was set as follows: 2% B, 1 min; 2%–100% B, 18.0 min; 100% B, 22.0 min; 100%–5% B, 22.1 min; 5% B, 25 min. QTRAP 6500+ mass spectrometer was operated in negative polarity mode with curtain gas of 35 psi, collision gas of medium, ion spray voltage of −4500V, temperature of 500°C, ion source gas of 1:55, and ion source gas of 2:55. Detection of the experimental samples using MRM were based on Novogene in-house database. Q3 (daughter) were used for the quantification. Q1 (parent ion), Q3, retention time, declustering potential and collision energy were used for metabolite identification. Data files generated by HPLC-MS/MS were processed with SCIEX OS (version 1.4) to integrate and correct the peaks. A total of 326 compounds were identified in the hemolymph samples. Metabolomics data analysis was then performed using MetaboAnalyst 3.0[43]

### RNA extraction and brain gene expression

Brain tissues of individual bees were collected using a dissecting microscope (Canon). Individual bee was fixed on beeswax using two insect needles through the thorax. After removing the head cuticle, whole brain was taken out on the glass slide placing on top of an ice pack. The hypopharygeal glands, salivary glands, three simple eyes, and two compound eyes were carefully removed in one drop of RNAlater (Thermo Fischer; Waltham, MA, USA). Dissected brains were kept frozen at −80°C. Total RNA was extracted from individual brains using the Quick-RNA MiniPrep kit (Zymo Research). RNA degradation and contamination were monitored on 1% agarose gels, and the purity was checked with the NanoPhotometer spectrophotometer (IMPLEN; CA, USA). RNA integrity was assessed using the RNA Nano 6000 Assay Kit of the Bioanalyzer 2100 system (Agilent Technologies, CA, USA).

RNA sequencing libraries were generated using NEBNext Ultra RNA Library Prep Kit for Illumina (New England BioLabs; Ispawich, USA) and index codes were added to attribute sequences to each sample. The clustering of the index-coded samples was performed on a cBot Cluster Generation System using TruSeq PE Cluster Kit v3-cBot-HS (Illumina), and the library preparations were then sequenced on an Illumina NovaSeq 6000 platform and 150 bp paired-end reads were generated. Sequencing quality of individual samples was assessed using FastQC v0.11.5 with default parameters. An index of the bee reference genome (Amel_HAv3.1) was built using HISAT2 v2.0.5[44] and the FastQC trimmed reads were then aligned to the built index using HISAT2 v2.1.0 with default parameters. StringTie v1.3.3[45] was then applied to assemble the obtained alignments in a BAM format into potential transcripts, using the GTF file of the honeybee genome downloaded from NCBI as a reference. Then we merged transcripts from all samples and examine how the merged-transcripts compare with the reference annotation. The gene and transcript abundances were estimated using merged-transcripts as reference.

Before differential gene expression, we first transformed the abundance into raw counts using scripts offered by StringTie. Count-level data underwent relative log expression (RLE) to estimate size factor and dispersion, then fit for each gene with fitType ‘local’, followed by Wald test to determine differential gene expression (DGE) between MF bees and bees mono-colonized with *Bifidobacterium asteroides* strain W8113 from “Bifido-1.4” using the R package DESeq2 v1.22.2[46]. Analysis of event-level differential splicing was performed using rMATS v4.0.2[47] based on the new merged-transcripts in StringTie as reference. An exon-based ratio metric, commonly defined as percent-spliced-in value, was employed to measure the alternative splicing events. The percent spliced in (PSI) value is calculated as follows:

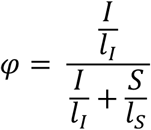

where S and I are the numbers of reads mapped to the junction supporting skipping and inclusion form, respectively. Effective length l is used for normalization. The PSI value were calculated for different classes of alternative splicing events, including skipped exon (SE), alternative 5’ splice site (A5SS), alternative 3’ splice site (A3SS), mutually exclusive exons (MXE), and retained introns (RI). Events with p < 0.05 were considered differentially spliced bwtween the two groups of bees.

### Statistical Analysis

Wilcoxon test was used to determine the significance of Bray-Curtis distance or the relative abundance of each microbiota taxon between each two bee groups and Kruskal-Wallis test was used for multi groups, and *P value* <0.05 indicating statistical significance. Box plots display first quartile, median, and third quartile. Mean values were noted in the beeswarm plots. The threshold for genome-wide significance was corrected for multiple testing with a weighted Bonferroni adjustment, with adjusted *P* <0.05 as significant. GLM and MLM are implemented for association tests.

## Results

### Gut communities are more different between genetically varied honeybees

*A. mellifera* belonging to four different subspecies, namely OH, AF, YF and SK, were sampled. They were imported into China in the 1980s, and were then kept in Jilin province for germplasm conservation. Metagenomic sequencing of gut homogenates from 57 individual bees from the four subspecies was performed, and bees from YK and SK were sampled from two independent hives each. We processed the whole gut for shotgun sequencing, thus simultaneously acquiring the genomic information of the host and microbial community. For each sample, 53–127 million pair-end reads (150 bp) were generated. First, in order to determine the genomic diversity of the hosts, the sequencing reads were mapped to the honeybee genome assembly (version Amel_HAv3.1). A total of 33–77% of the reads were mapped to the honeybee genome, indicating a 13–57× coverage of the honey bee genome, and 2,255,909 sites were identified as polymorphic (Figure S1a). An evolutionary tree of *A. mellifera* inferred from all single-nucleotide polymorphism (SNP) demonstrated clear clustering of the four groups, and the OH bees were more distantly related to the other three subspecies (Fig. 1a). Likewise, ADMIXTURE analysis of genetic co-ancestry also partitioned the data population into the four defined groups when the number of populations was set at *K*=4 (Fig. 1b, Figure S1b). Different subspecies had an average pairwise *F*_ST_ (allelic fixation index) of 0.08–0.11, which was consistent with a previous analysis of *A. mellifera* subspecies[48]. These results supported the designation of the subspecies with distinct genetic backgrounds.

**Fig. 1.**
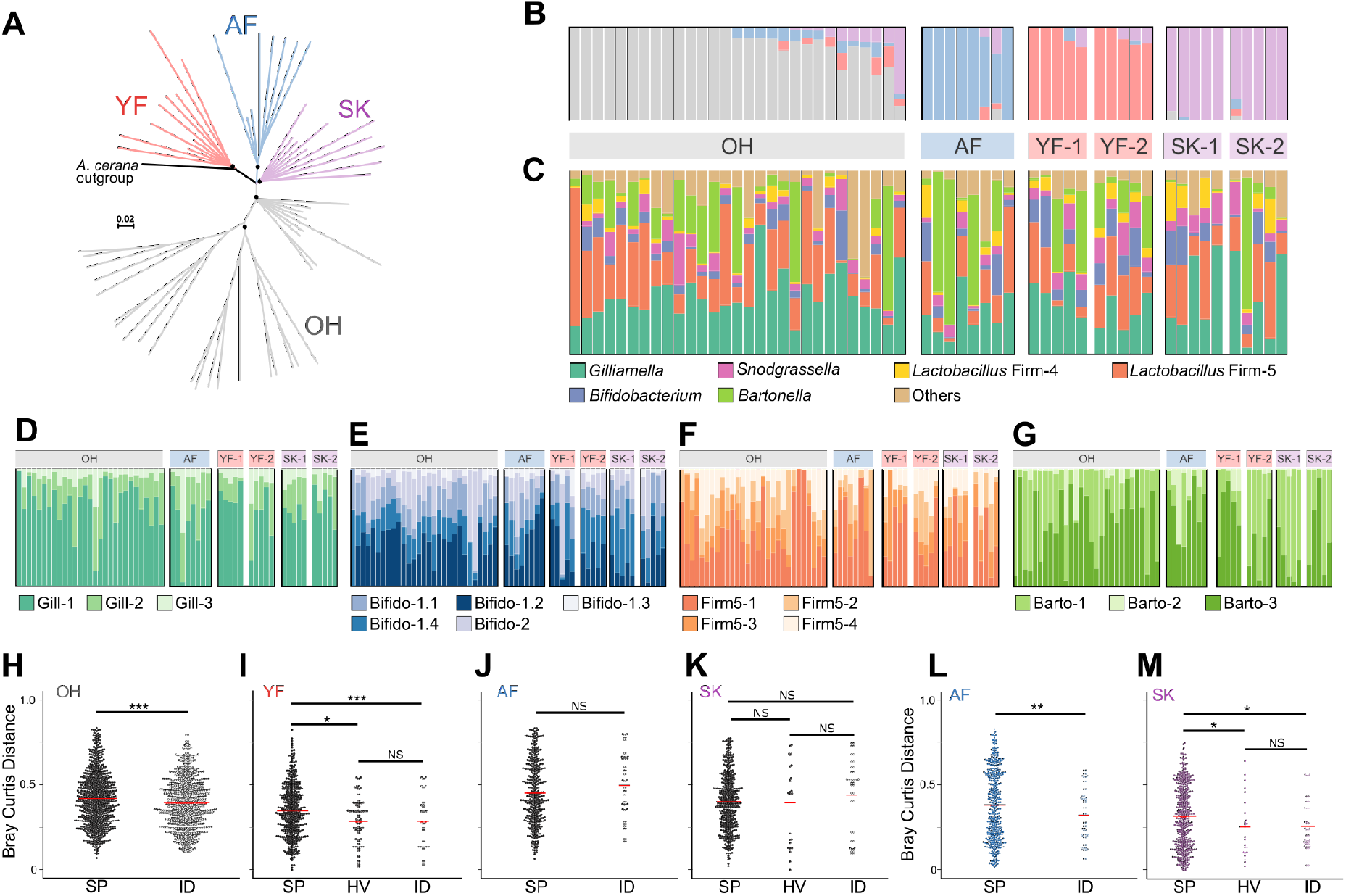
Gut microbiota compositions differ across genetically varied honeybees at both phylotype- and SDP-level. **a**, Neighbor-joining tree constructed from allele-sharing distances between honeybee subspecies. Nodes with 100% support are marked with dots. The scale bar represents raw genetic distance per variable site. **b**, ADMIXTURE analysis showing clustering of bee samples into four groups (*K*=4). One colony each of OH (n=29) and AF bees (n=8), and two colonies each of YF (n=10) and SK (n=10) bees were sampled for the metagenomic analysis. Each bar represents one bee individual. **c**, Relative abundance of phylotypes in the guts of bee individuals from different subspecies. (**d**–**g**) Relative abundance of sequence-discrete populations (SDPs) for four core gut members: *Gilliamella* (**d**), *Bifidobacterium* (**e**), *Lactobacillus* Firm-5 (**f**), and *Bartonella* (**g**). Dissimilarity of phylotype- (**h**–**k**) and SDP-level (**l**, **m**) gut composition profiles between paired samples as measured by Bray-Curtis distance. The distances are separated by the category of comparison: comparison between bees from different subspecies (SP), between bees from different hives of the same subspecies (HV), between individuals from the same colony (ID) (NS, not significant; *, p<0.05; **, p<0.01, ***, p<0.001; Wilcoxon test).

We then assessed the composition of the gut community using the MIDAS pipeline with a custom database[33]. To construct a better reference database for the analyses of strain-level genomic variation, we isolated 116 bacterial strains from the gut homogenates of *A. mellifera*, and a new genomic database was generated using 405 bacterial genomes from both honey and bumble bee gut isolates (Figure S2, Additional file 1: Dataset S1). For all samples, an average of 26% of reads was mapped to the bacterial database. The relative abundances of bacterial phylotypes and SDP were estimated using MIDAS, which maps reads to a panel of 15 selected marker genes of the genomes (see Methods). Although many reads were mapped to the host genome, the accumulative curves of the observed SDPs begun to plateau, indicating that the microbial dataset was adequate for the diversity analyses (Figure S3). Consistent with previous 16S rRNA- and metagenome-based studies[19, 49], all individual gut communities were dominated by the five core bacterial species. *Gilliamella* and *Lactobacillus* Firm-5 were the most abundant members, whereas some AF and OH bees had a higher fraction of *Bartonella* (Fig. 1c). Although it was suggested that wintering bees possess more *Bartonella*[50], this study was performed in July 2018, and all individuals were sampled on exactly Day 15 following the emergence, suggesting it is not a seasonal or age effect. In addition to the core members, the other non-core species, including *Frischella perrara*, *Lactobacillus kunkeei*, and *Commensalibacter* spp., were detected in variable amounts. Since the honeybee gut community had a low complexity at the phylotype level, the SDP-level profiles of different bee subspecies were then compared. First, all SDPs from each phylotype co-occurred in individual bees. While the relative abundance of different phylotypes was not obviously different, the SDP-level profiles were more variable between individuals with different genetic backgrounds. For example, although the frequency of *Bifidobacterium* was not different among subspecies of bees, OH and AF had more strains from the Bifido-1.2 SDP, while Bifido-1.4 was more abundant in YF and SK bees (Fig. 1e, Figure S1c). For *Gilliamella*, Gilli-1 was the predominant SDP, while Gilli-3 was only present in very small amounts in all bees and the frequencies were not associated with host genetics (Fig. 1d, Figure S1c).

To compare the bacterial communities of different subspecies of bees, we examined the variability within and across host groups by testing the Bray-Curtis dissimilarity between individuals from i) different subspecies (SP); ii) bees from different hives but belonging to the same subspecies (HV); and iii) individuals from the same hive (ID). At the phylotype-level, the dissimilarities were significantly higher for the comparisons between genetically varied bees (SP) than those for bees from the same subspecies (HV and ID; Fig. 1h, i), suggesting an effect of host genotype. However, the same category comparisons for AF and SK bees were not significantly varied (Fig. 1j, k), possibly due to the low resolution and the few phylotypes present in the bee gut. Therefore, we further calculated the distances at the level of SDP, and the dissimilarities between different subspecies of individuals for both AF and SK bees were significantly higher than those of bees with similar genetic backgrounds (Fig. 1l, m). Notably, for YF and SK bees, the inter-hive distances were not different from those of individuals from the same hive, suggesting that the hive effects were limited here. In summary, our results showed that the gut community compositions of bees with a varied genetic background were more different than those of individuals from the same bee subspecies, indicating that the host genetic variation is associated with the gut microbiota profiles.

### Host genotype determines the composition of passaged gut community

The gut microbiota of honeybees are acquired via social interactions with other workers, but not through a maternal vertical transmission[51]. Thus, the gut microbiota is transmitted in the colony between batches of siblings from the same queen. To determine if the host genotype impacts the socially transmitted microbiota, we started two lines of colonies (three replicate hives each) headed by OH virgin queens instrumentally inseminated with semen from single drones of AF or YF (Fig. 2a). Then colonies were initiated by the inseminated queens, together with ~300 founding workers (O-A’ and O-Y’) randomly sampled from one hive without control of host genetic background, who fed the hybrids at the beginning (O-A and O-Y). Thus, the gut microbiota of the hybrids must have been derived from the founding workers, and then transmitted among batches of hybrids within the colonies. Three different batches of newly emerged hybrid adults were marked with color paints, and they were sampled when they were exactly 15 days old. It is noteworthy that, before the third batch of bees started to emerge, all initial founding workers had died.

**Fig. 2.**
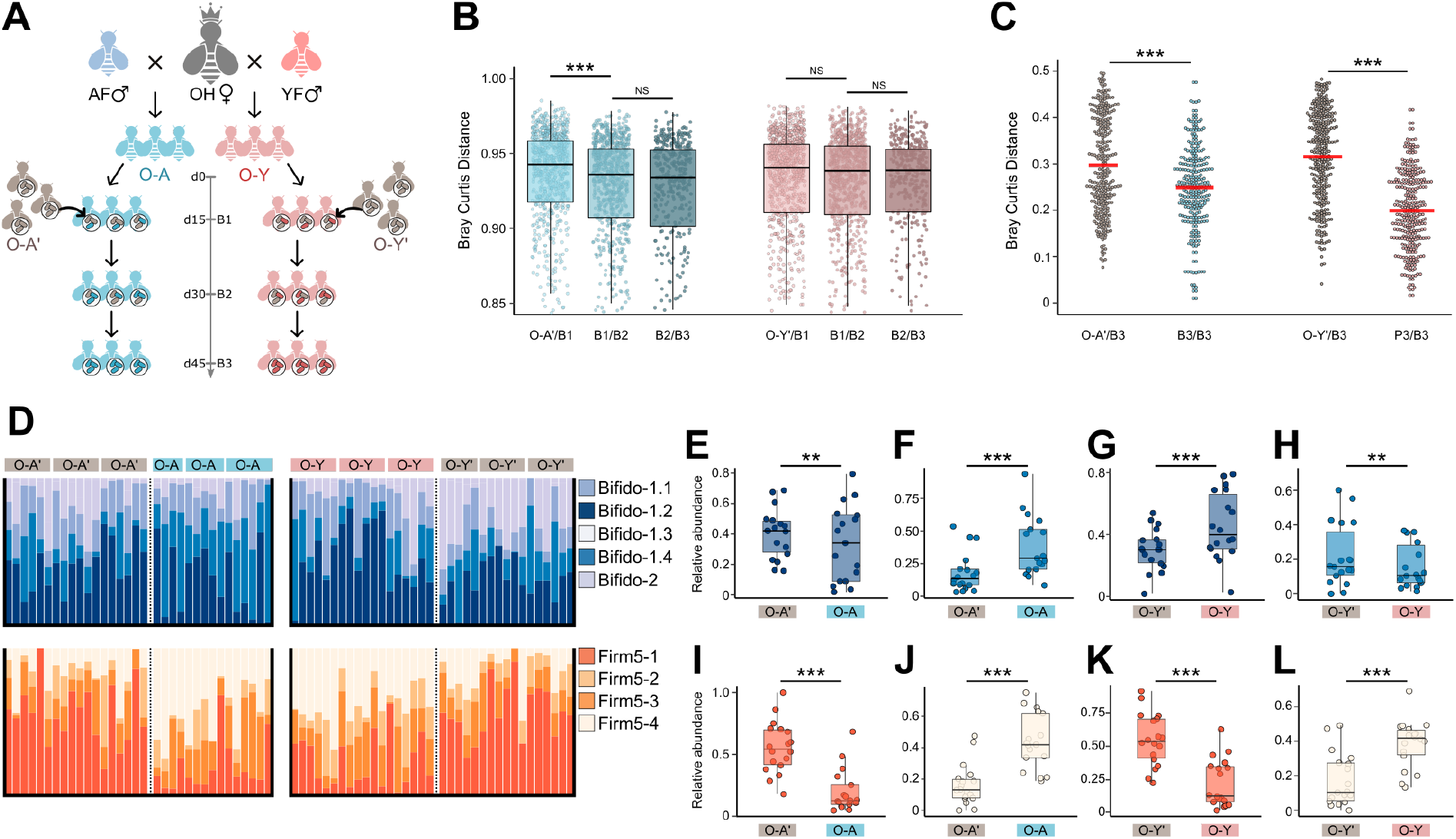
Gut community shifts during the successive passage within the colony. (**a**) Study design for serial transmission of gut microbiota in two lines of hybrid bees generated by artificially insemination. We sampled guts from three batches of hybrid bees (B1–B3) during the passage and from the founding workers that initiate the colony (O-A’, O-Y’). (**b**) Bray-Curtis distance of the gut communities between bees from different batches at the OTU-level of 16S rRNA sequences. (**c**) Bray-Curtis distance of the gut communities at the SDP-level by metagenomic sequencing. (**d**) Relative abundance profiles of SDPs from *Bifidobacterium* and *Lactobacillus* Firm-5 for the founding workers (O-A’, O-Y’) and the B3 batch of individuals (O-A, O-Y). (**e**–**l**) Comparison of the relative abundances of the SDPs of *Bifidobacterium* Bifido-1.2 (**e**, **g**), Bifido-1.4 (**f**, **h**), *Lactobacillus* Firm5-1 (**i**, **k**), and Firm 5-4 (**j**, **l**) between the founding workers and the B3 individuals. Wilcoxon test was used to compare the average of Bray-Curtis distance or the relative abundance of each microbiota taxon between each two bee groups, and *P value* <0.05 indicating statistical significance. (NS, not significant; **, p < 0.01; ***, p < 0.001; Wilcoxon test).

The genetic backgrounds of the founding workers were genetically different from the hybrid bees (Figure S4a). Next, the impact of host genotypes on bacterial community transmission was measured by 16S rRNA sequencing of the gut microbiota of the founding bees, together with the three batches of hybrid workers (B1–B3). By testing OTU-level Bray-Curtis dissimilarity between adjacent time points for each batch of individuals, it was found that gut communities of hybrids with more similar genetic backgrounds (B1/B2 and B2/B3) exhibited similar features over time (Fig. 2b). However, gut communities of B1 markedly changed from those of the founding bees of O-A’. While the gut community did not shift between O-Y and O-Y’ bees at the phylotype level, we next determine the fractions of all SDPs in the founding bees and the B3 batch (Figure S4c). Overall, the compositions were much more similar within B3 than those between B3 and founding workers at the SDP level (Fig. 3c). While the relative abundances of most SDPs were not significantly different, the SDPs Bifido-1.2 and Bifido-1.4 were differentially distributed in the genetically varied hosts (Fig. 2d). Interestingly, Bifido-1.2 was enriched in the O-A’ founding workers, as compared to O-A, however, it was more abundant in the B3 of O-Y bees (Fig. 2e, g). By contrast, Bifido-1.4 were more abundant in O-A, but were decreased in O-Y, as compared with the founding workers (Fig. 2f, h). These strongly suggested that the host genetics showed different selection powers upon the bacterial SDPs. For *Lactobacillus* Firm-5, all founding workers had a higher fraction of the Firm 5-1 cluster, whereas the B3 bees harbored more Firm 5-4 in the gut (Fig. 2i–l). Altogether, our results indicated that some SDPs shifted in relative abundances in hosts with differential genetic backgrounds during transmission, and the host genotype could shape the pattern of gut microbiota transmission within the colony.

**Fig. 3.**
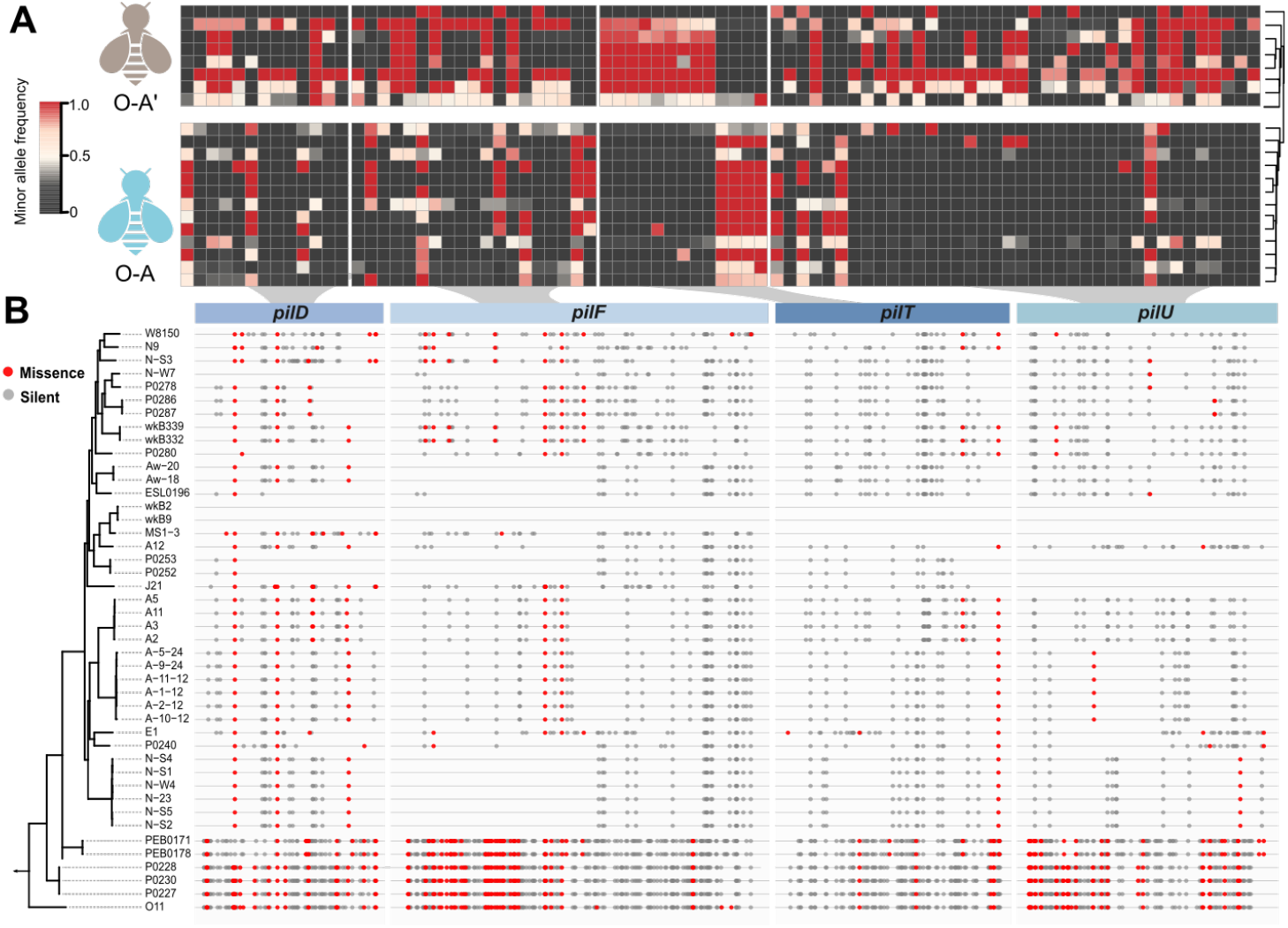
SNPs of Type IV pili component genes of *Snodgrassella alvi* are differentially distributed between the founding workers and the B3 bees in the colony. (**a**) A heatmap showing the minor allele frequency for missense SNPs that are significantly different in founding workers (O-A’) and the B3 batch of bees (O-A). Each row represents one bee metagenomic sample, each column is one site in the T4P genes. The tree on the right illustrates a dendrogram of clustering (Ward’s method). (**b**) Whole-genome phylogenetic tree of isolated *Snodgrassella* strains (Additional file 1: Dataset S1) using the maximum-likelihood algorithm based on the concatenation of core protein sequences. The lines aligned to tree leaves represent corresponding gene sequences with missense (red dot) and silent (grey dot) SNPs indicated.

### Biased SNP distribution in Type IV pili (T4P) structural component-coding genes underlying the strain-level difference in *Snodgrassella*

It has been shown that strains of the same SDP from the bee gut are genetically divergent, and the strain-level profiles can be different among individuals from the same colony[19]. Unlike other core members, *Snodgrassella* possess only one SDP, and they specialize in colonizing the hindgut epithelium, suggesting its relatively close interactions with the host[52]. To identify if hosts with different genetic backgrounds have characteristic strain compositions, we analyzed the differences in genome SNP distribution of *Snodgrassella* strains between the founding workers and the B3 individuals. Herein, the distribution of the minor allele frequency per genomic site was used to indicate the strain variations in different hosts. An examination of the SNP distributions of protein-coding genes of *Snodgrassella* revealed 1,547 genes with a valid coverage in 52 bee individuals. Among these genes, 1,436 genes possessing sites with significantly differentiated allele frequencies were identified between the two groups of bees (Mann-Whitney test, p < 0.05). Notably, four genes encoding the T4P harbored the most differentiated SNP distributions with significant enrichment of group biased SNPs between O-A and O-A’ bees, while the coverage for O-Y group was not sufficient for the downstream analysis (Fig. 3a, Additional file 2: Dataset S2). Firstly, with the reference strain of wkB2, we identified 1,017 SNPs in the *pilD* (prepilin peptidase), *pilF* (fimbrial biogenesis protein), *pilT* (pilus retraction/twitch motility motor), and *pilU* (prepilin peptidase) genes in the genomes of *Snodgrassella* isolates, and there were generally more SNPs (both missense and silent) in phylogenetically distant strains (e.g., strains PEB0171 and P0228; Fig. 3b). The heatmap presented all missense SNP sites with significantly biased distributions between the O-A and O-A’ groups (Mann-Whitney test, p < 0.05), and the dendrogram based on SNP frequencies exhibited two different clustering groups, according to the host genotype (Fig. 3a). These indicated that: i) *Snodgrassella* strains exhibited a markedly different enrichment of SNPs in the T4P genes; ii) a specific set of strains were found to be correlated with genetically varied bee hosts. A genome-wide Tn-seq analysis has documented that the T4P genes of *Snodgrassella* are beneficial for gut colonization, and are potential determinants of host specificity[52]. These indicated that the host genotype may have a strong effect on the strain-level composition of gut bacteria, and that the T4P of bacteria play vital roles in microbiota-host interactions[53].

### GWAS revealed that the relative abundance of *Bifidobacterium* is associated with host genetic variants

Since a correlation was observed between the host genotype and the gut composition, the heritability of each taxa was then estimated, and GWAS was performed to identify which host gene was most associated. Both the host genotype and gut community composition data of 125 individuals (including 57 pure bees, 33 hybrid bees, and 35 founding workers) were included in this analysis. The proportions of six core bacterial phylotypes and 17 SDP-level compositions were treated as individual traits, and 2,861,994 informative SNPs spaced throughout the honeybee genome were included (Figure S5a). We used both a mixed-model algorithm and an interactive usage of fixed and random effect models to correct the population structure for GWAS. The threshold for genome-wide significance was corrected for multiple testing with a weighted Bonferroni adjustment, according to the simulation and permutations, as previously described[54]. While most of the associations did not reach the study-wide significance, the compositions of the Gilli-2 and Bifido-1.4 SDPs were found to be significantly associated with host genotype (Figure S5a). Heritability analysis revealed that both SDPs showed a relatively higher heritability (Figure S5b).

GWAS using both GLM and MLM revealed that the SNPs with the strongest association to the relative abundance of the Bifido-1. 4 SDP lies within the *gluR-B* gene located on chromosome 13 (p < 0.05; Fig. 4a and 4b). GluR-B is a metabotropic glutamate receptor (mGluR) involved in the regulation of glutamatergic synapses, and is specifically expressed in the honeybee brain[55]. A close observation showed that the *gluR-B* gene was enriched by 112 SNPs highly associated with the Bifido-1.4 SDP (Fig. 4c). Correspondingly, bee individuals carrying the TC/CC allele have higher levels of Bifido-1.4 than those carrying the TT allele at the locus with the strongest association (chr13-7527228; Fig. 4d). In addition, the relative abundance of Gilli-2 in the gut is correlated with the SNPs from multiple genes encoding inositol-pentakisphospate 2 kinase, carcoxypeptidase Q and poly(rC)-binding protein within chromosome 12 (Figure S5).

**Fig. 4.**
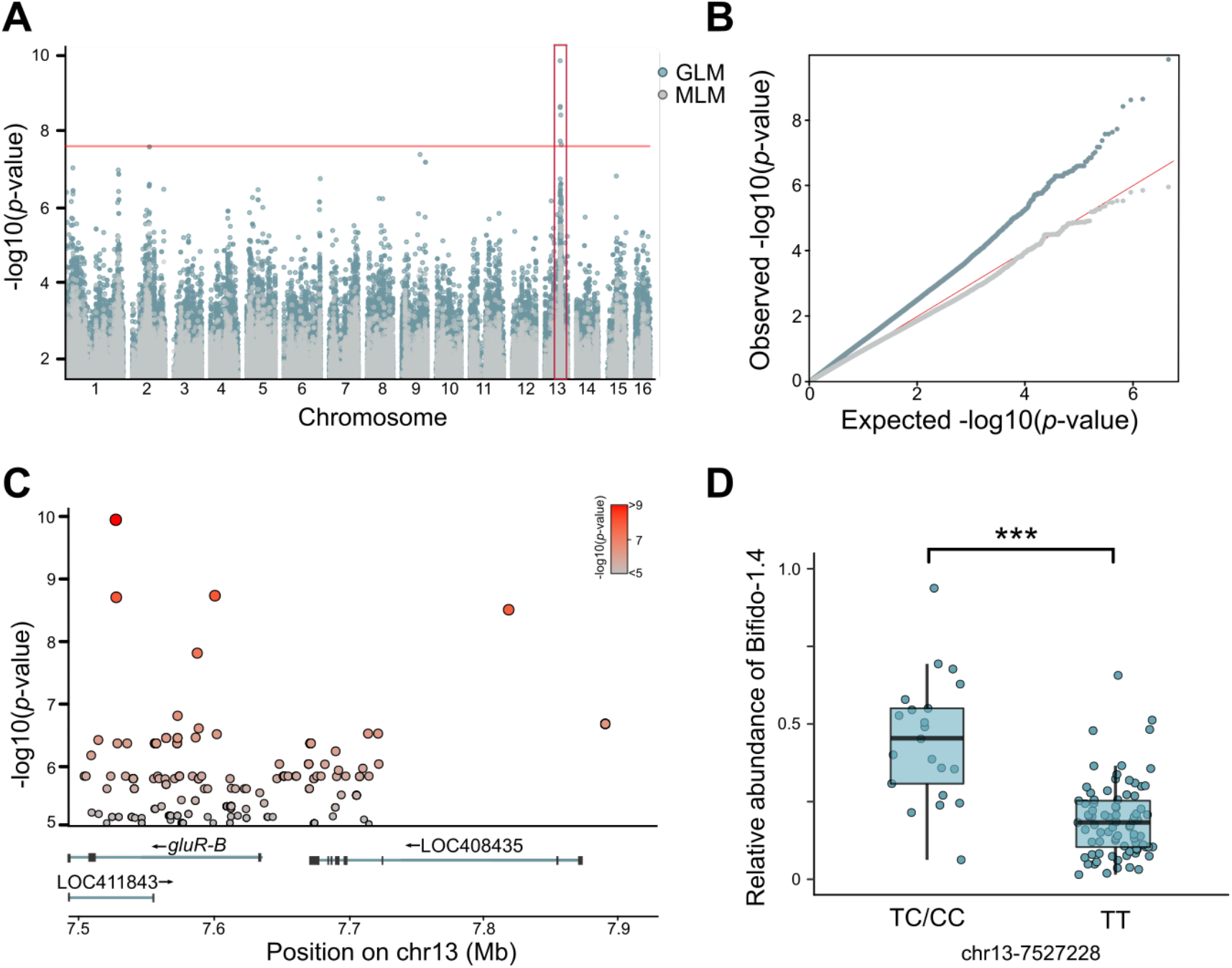
Relative abundances of the heritable *Bifidobacterium* SDP Bifido-1.4 are associated with genetic variants in the genomic locus containing the metatropic glutamate receptor gene. (**a**) Genome-wide Manhattan plot: each dot represents the −log of the p value for the association of the SNP with the relative abundance of the Bifido-1.4 SDP. The red box highlights the associated locus on chromosome 13 containing the gene *gluR-B*. The threshold for genome-wide significance was corrected for multiple testing with a weighted Bonferroni adjustment. The threshold value was set at - log(p) > 2e-8. We used both General Linear Model (GLM) and Mixed Linear Model (MLM) for the association tests (**b**) Quantile-quantile plot showing deviation from the expected distribution of p values. The diagonal (red) line represents the expected distribution. (**c**) Close-up plots of ~0.4 Mb window around the SNPs with the highest associations. The coloring of each circle is proportional to the significance. The exon intron architecture of the *gluR-B* gene and two neighboring genes (LOC408435, LOC422843) are shown at the bottom. (**d**) Relative abundance of the Bifido-1.4 SDP in bees with different genotypes at the *gluR-B*-associated SNP (chr13-7527228; p < 0.001, Wilcoxon test).

### *Bifidobacterium* alters gene expression and alternative splicing of *gluR-B* by elevating the GABA level in brain

The honeybee mGluR is a G-protein coupled receptor (GPCR), which shares sequence similarity with the B-type gamma-aminobutyric acid (GABAB) receptors, the calcium sensing receptors, and some pheromone receptors[56]. The mGluR and GABA receptors mutually modulates signal transduction, and its expression is controlled by both glutamate and GABA in the brain[57]. GABA is an inhibitory neurotransmitter found at high concentrations in the honeybee brain, which has been implicated in several honeybee behaviors, including odor coding, learning and memory, as well as locomotion control[58, 59]. The altered gut and hemolymph metabolome have been associated with the honeybee microbiome; specifically, glutamic acid is enriched both in the gut and the hemolymph of bees with a normal gut community[24]. Since the significant association between GluR-B and the level of gut *Bifidobacterium* were identified, whether the concentration of honeybee brain GABA is affected by the colonization of Bifido-1.4 was next explored. Targeted metabolomics revealed that the GABA concentration in brains of bees mono-colonized (MC) with Bifido-1.4 was ~1.5 times higher than that of MF (Fig. 5a).

**Fig. 5.**
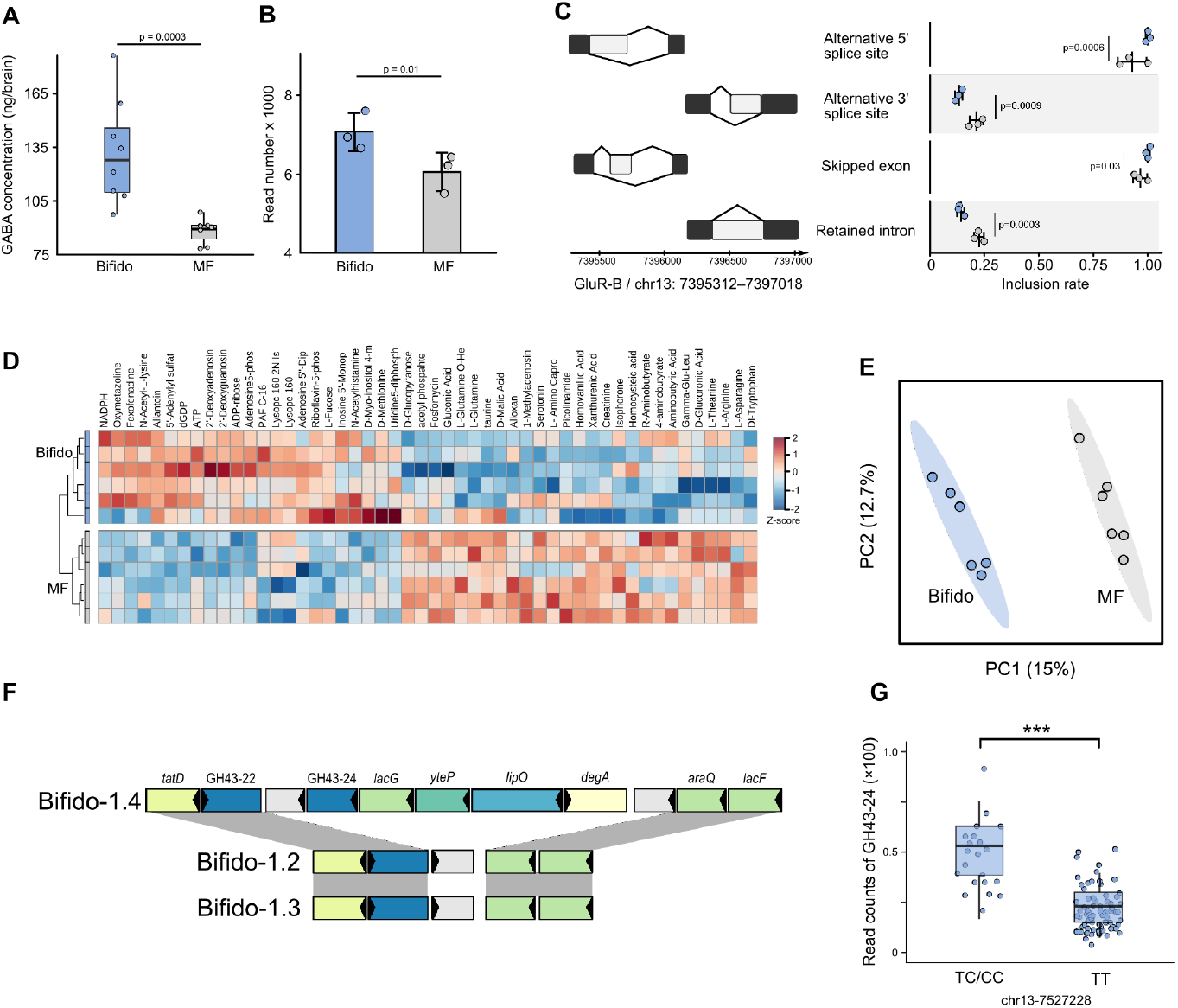
*Bifidobacterium* impacts gene expression and GABA level in the brain, and affects the hemolymph metabolome of honeybees. (**a**) Targeted metabolomics indicates that the GABA concentrations are increased in the brain of Bifido-1.4 mono-inoculated honeybees than the microbiota-free (MF) bees. (**b**) Relative expression levels of *gluR-B* gene in brains of MF and Bifido-1.4 inoculated bees. (**c**) Differential splicing events of *gluR-B* gene in the brains of MF and Bifido-1.4 inoculated bees. Benjamini-Hochberg corrected p values are shown. (**d**) Heatmaps of differentially abundant metabolites identified by a quasi-targeted metabolomics of hemolymph. Colors indicate the Z-score based on the normalized relative concentration of each metabolite. The tree on the left illustrates a dendrogram of clustering (Ward’s method). (**e**) Partial least squares discriminant analysis based on 328 metabolites (Additional file 3: Dataset S3) detected from the hemolymph of MF (n = 6) and Bifido-1.4 inoculated bees (n = 6) showing different clustering groups (95% confidence regions). (**f**) Syntenic loci of the PUL in *B. asteroides* strains from different SDPs. Homologous genes are connected by gray bars. (**g**) Boxplots of the read counts for GH43-24 specific to Bifido-1.4 in each genotype at the *gluR-B*-associated SNP (chr13-7527228).

Given evidence that *gluR-B* is preferentially expressed in the brains, we investigated whether the colonization of *Bifidobacterium* altered the expression level. RNA sequencing analysis revealed that *gluR-B* was downregulated in MF bees (Fig. 5b, Additional file 3: Dataset S3), suggesting that the glutamate receptor might also be regulated by GABA in the brain[60]. In addition, we detected the alternative splicing (AS) events in *gluR-B*, and rMATS analysis[47] indicated that *gluR-B* exhibited differential patterns of AS events between MF and MC bees. Bees colonized with Bifido-1.4 showed increased inclusion rates of 5’ alternative start site and skipped exon events in *gluR-B*, while the event frequencies of 3’ alternative start sites and retained introns were lower in MF bees (Fig. 5c). These results suggested that gut microbes not only regulate gene expression in the bee brain, but also affect AS, which regulates the production of specific isoforms of the mGluR gene.

### *Bifidobacterium* produces discrete metabolite profiles and gene abundance differences are potential drivers in strain transmission

Brain gene expression, slicing and neuronal functions can be regulated by the gut microbial metabolism[61], and the altered gut metabolome has been observed between MF bees and bees colonized with a conventional gut microbiota[24]. Untargeted metabolomics analysis of hemolymph from MF and MC bees was performed, and 128 out of 326 detected metabolites were significantly different (Additional file 4: Dataset S4). Significant differences were identified in many amino acids between the two groups, similar to previous reports, when comparing MF and conventional bees[24]. Notably, both GABA and taurine were significantly lower in MC bees (Fig. 5d), which was consistent with our observation in the brain. Taurine is a weak GABAA receptor agonist correlated to behavioral disorder in humans and mice[62]. This suggests that *Bifidobacterium* may impact endocrine inhibitory GABA signaling. The partial least squares-discriminant analysis showed that *Bifidobacterium* clearly altered the hemolymph metabolome (Fig. 5e).

Finally, we sought to identify gene presence/absence signatures that may explain the inheritance patterns of different *Bifidobacterium* SDPs and their ability to alter brain functions. We wonder why strains of the Bifido-1.4 SDP are preferentially transmitted and hypothesized that these strains possess specific genes that are implicated in their interactions with the host. It has been documented that *Bifidobacterium* are the major degraders of diet hemicellulose, and the abilities of individual strains vary[32]. We therefore searched for genes that are present in Bifido-1.4, but absent in the other SDPs of *Bifidobacterium*. A total of seven genes that are specific to Bifido-1.4 strains were identified, and they were located in a single genomic region of the reference genomes (Fig. 5f). This region contained two carbohydrate-active enzymes (CAZyme) of GH43-22, together with *lacG* encoding phospho-beta-galactosidase, the LacI-family transcriptional regulator *araQ*, the helix-turn-helix type transcriptional regulator *degA* and a multiple-sugar transport system permease *yteP*. They form a typical structure of the CAZyme gene cluster, which may enable them to specialize in the breakdown of dietary fiber. To confirm if host genetics are associated with the functional capacity of the gut bacteria, the read counts of the GH43-22 gene were compared in bees showing different alleles. Our metagenomic analysis revealed an enrichment of the glycoside hydrolase GH43-22 in the guts of individuals with TC/CC at the locus chr13-7527228, suggesting that the bee genotype is associated with an abundance in polysaccharide-degrading gut microbes.

## Discussion

In the present study, the socially transmitted gut microbiota of pure-bred and hybrid honeybees were used to elucidate the consequences of host genetic divergence in the composition of symbionts. Our metagenomic analysis characterizing the microbial structure at different taxonomic levels represents strong evidence that specific gut members are affected by the host genetic factor during the transmission. Moreover, bee individuals were genotyped simultaneously with the characterization of the microbiome, allowing for tests of the association between host SNPs and microbiome traits. Finally, we focused on the heritable gut member, *Bifidobacterium* sp. Bifido-1.4, and its colonization altered the brain neurotransmitter and gene expression patterns, which might be associated with a unique PUL-like region in the genome.

It has been well documented that the honeybee gut microbiota are dominated by limited numbers of bacterial phylotypes, commonly with species from the *Gilliamella*, *Snodgrassella*, *Lactobacillus*, *Bifidobacterium* and *Bartonella* genera. Therefore, it is not surprising that the variation was not obvious among individuals at the phylotype-level (Fig. 1). Although the core gut members have been observed in bees across studies, even in samples from different countries, gut microbiota can differ markedly in diversity across adult bees[17]. Several factors have been found to be influential in the composition of the bee gut community, such as the regional floral diversity, seasonal shift, age and bee caste[27, 63]. Nevertheless, the environmental factors may all contribute to diet preference and nutritional status, which are the main drivers of community variance. However, certain studies have also reported an intercolonial difference among individuals belonging to the same caste and sampled at the same age. Specifically, high levels of strain-level diversity exist within all major phylotypes in the honeybee gut. By targeting protein-coding gene markers of *Gilliamella* and *Snodgrassella*, it was found that strain compositions of individuals from the same hive can vary dramatically, and some strains are specific to only one host species[22, 64]. Such strain-level diversity was also recently appreciated by community-wide shotgun sequencing[20]. Since all studies have focused on colonies naturally headed by hyperpolyandrous queens, we hypothesized that the diversity in the bee gut microbiota was shaped by both environmental factors and host genetics. Our findings clearly showed that the abundance in both phylotypes and SDPs were more correlated within bees from the same subspecies, indicating the host genetic interactions with specific gut members. In humans, comparative analyses revealed that family members have more similar microbiota, partly due to shared environmental influences[65, 66]. Although we sampled bees from outdoor hives, we controlled individual age and sampled simultaneously from multiple colonies reared at the same apiary, to minimize factors other than host genetics. While identical environmental conditions cannot be guaranteed for these colonies (e.g., possibly a bias for flower preference), no bias in the flower preference of honeybees was identified in host genetics[67].

In bee colonies, the gut symbiotic bacteria are transmitted between generations of siblings through social contact, and thus the core lineages of gut bacteria show phylogenies matching those of the hosts, highlighting codiversification along the evolutionary history of symbiosis. Mounting evidence have shown codiversification via a long-term vertical association in many animal groups, and host filtering could be a major driver of “phylosymbiosis”[1]. In our microbiota passaging approach in two independent lines of hybrid bee populations, a considerable shift in community composition was found during the transmission. Notably, the highest divergence was observed between the founding workers and B1, indicating that host genotype shapes the gut composition early in passaging. Although it was found that all SDPs from the core phylotypes can be inherited by the hybrid bees during the consecutive transmission within the hive, the effect of the host differed widely on even closely associated discrete bacterial populations from the same phylotype. For two SPDs from *Lactobacillus* Firm-5, the host filtering processes were similar in O-A and O-Y hybrids. By contrast, the relative abundance of *Bifidobacterium* Bifido1.2 was decreased in O-A but increased in O-Y bees (Fig. 2). This shows that genetically varied hosts permit the colonization of specific sets of bacteria, strongly suggesting that the effect of the genotype is driving the microbiota structure. While we observed an obvious shift in gut community between the founding bees and the first batch of O-A, the microbiota was not significantly altered during transmission in O-Y (Fig. 2b). This suggests that the effects of genetic variance among hosts could be different, and interestingly, the divergence of host genetics in the O-A group was lower than that among O-Y hybrids (Figure S4a), which might partly explain the observed filtering process. When the living environments and microbial populations are controlled, the influences of host genetics on microbial compositions have also been detected in other animals and plants[68, 69]. However, the composition of resultant output microbiota is always determined by the input species[70]. In our transfer experiment, the output microbiomes were compared within two parallel lines with a nearly identical starting community carried by the same batch of founding workers; however, they were passaged in genetically different hosts, allowing us to overcome the legacy effects (Fig. 2).

So far, only one SDP from *Snodgrassella alvi* was identified, but a markedly high level of strain diversity was observed based on the proportion of polymorphic sites in core gene sequences[19]. For bees from the same colony, it was found that strains can be dominant in one individual and totally absent from another[64]. Although the GWAS associated host genetic loci to the overall microbiome divergence (beta-diversity) or the abundances of several taxa, few studies have focused on the strain composition, which is mostly due to the markedly high level of diversity at this level. The presence of only one lineage of *Snodgrassella* with genetically divergent strains enables the fine-scale analysis of the effect of the host on strain stability. It was found herein that strains are subject to the selection of host genotype, and different strains are enriched in genetically varied hosts during the social transmission. This suggests that bees with different genetics have an elaborate mechanism to ensure a specificity of the association, which might be achieved by signal recognition and secretion of antibacterial agents, as found in the symbioses of microbes and both plants and animals [71, 72]. Interestingly, it was found that the distribution of SNPs in T4P genes were significantly biased between genetically different hosts. T4P encodes membrane-associated transporter complex, which is important for biofilm formation and cell adhesion and has been identified as significant for the survival of pathogenic bacteria in eukaryotic hosts[73]. *Snodgrassella* forms a layer attached to the inner gut wall, which is vital for the maintenance of the gut microenvironment[24]. *Snodgrassella* possesses all core components of T4P in the genome, which are used to facilitate colonization *in vivo*[52]. Specifically, the ability of surface biofilm formation of structural mutant *pilF*^−^ was significantly reduced, and the SNP distribution of *pilF* was found to be obviously affected by the host genotype. Another major pilin subunit, *pilE*, which is beneficial for colonization, is exclusive to strains from honeybee, suggesting that T4P is a decisive factor in host adaptation. T4P are multicomponent transporters for the transfer of proteins and DNA into target cells, and are critical for the host specificity of most bacterial pathogens by mediating the adhesion and invasion processes[74, 75]. Our findings illustrated that T4P might provide selective advantages for different strains during the transmission in hosts with various genotypes, specifically for *Snodgrassella* colonizing the gut epithelium.

In our dataset, the Bifido-1.4 SDP from *Bifidobacterium* exhibited the most significant association with the host *gluR-B* gene locus and is also a highly heritable taxon. *Bifidobacterium* is also heritable in the human TwinsUK population, the HMP and mouse models[7, 76–78], which implies that *Bifidobacterium* are a group of symbionts critical for the physiology or metabolism of a variety of animal hosts. Indeed, it has been documented that *Bifidobacterium* are the principal polysaccharide degraders for bees, and not all members of SDPs are capable of digesting hemicellulose in diet pollen. Here, Bifido-1.4 with a strong signal in the GWAS are capable of degrading hemicellulose *in vivo* and possess an abundant repertoire of carbohydrate-active enzymes[32], highlighting that the diet interaction is fundamental for the host-gut microbe association. Indeed, in human intestines, the correlation between *Bifidobacterium* and SNPs near the *LCT* gene associated with lactase nonpersistence has been replicated across several populations[11]. This is due to the presence of lactose consumed by lactase nonpersister in the large intestine, which can stimulate the proliferation of *Bifidobacterium*[5]. While the specific strains of *Bifidobacterium* that are implicated in the association with the host genotype of lactose nonpersister were not determined in humans, our analysis specifically illustrated an association between host genetics and strains with more abundant glycoside hydrolase genes (GHs) that assist polysaccharide digestion in the bee gut. To characterize the functional significance of strain inheritance events, we examined genes only present in strains of Bifido-1.4 and not in other SDPs of *Bifidobacterium*. Bifido-1.2 and −1.4 both have an enrichment of GHs forming PUL-like regions in the genome; yet only one is different between these two SDPs, which might provide a selective advantage in colonization[79].

The sites significantly associated with Bifido-1.4 are located in *gluR-B*, which is a member of GPC mGluRs. Two subtypes of mGluRs have been characterized in honeybees, with only *gluR-B* expressed at high levels in the Kenyon cells within the mushroom body[55]. While they are both glutamate receptors, their overlapping expression in the brain suggests that they interact to modulate the glutamatergic neurotransmission and may also respond to different levels of GABA[80], one of the most abundant neurotransmitters in adult bee brains[81, 82]. Although the host genotype has not been linked to the GABA concentration in the brain, an altered expression level of *gluR-B* was observed with a higher level of GABA (Fig. 5). Taken together, these results point to an interaction between the *Bifidobacterium* and host brain physiology. The causality of the association between the relative abundance of the members of Bifido-1.4 and the host phenotype of brain neurotransmitter was tested experimentally by mono-colonization into MF bees. Mono-associated bees showed an increased level of GABA in the brain, as compared to their MF counterparts. Moreover, both the gene expression and the AS events of *gluR-B* are affected by the gut microbe, which can regulate the production of specific isoforms of genes implicated in host phenotypes. Gut bacteria can impact gene expression and host behavior through the production of neuroactive small molecules. Concentrations of several neuroactive metabolites, including serotonin, taurine, and aminobutyric acid in the circulation were regulated by *Bifidobacterium*.

Interestingly, a recent study has linked the human autism spectrum disorder to the microbial metabolism of 5-aminovaleric acid, a weak GABA_A_ receptor agonist[62], which is one of the most elevated metabolites in gnotobiotic bees with conventional microbiota [24]. These suggest that honeybee gut microbiota may regulate host phenotypes via the microbial metabolism, implying a gut-brain connection that contributes to impaired behaviors that share common molecular mechanisms with those of humans[83, 84].

Honeybees, as effective pollinators, are instrumental in the production of foods all over the world. Recent decline in the honeybee population has been a major threat to the balance of the global ecosystem. It has been shown that host genetic variation drives the honeybee phenotype, particularly the host nutritional status, colony health and productivity[85–87]. Our data illustrated that the socially transmitted gut microbiota assembled at emergence are clearly influenced by the host genotype. Given the evidence that bee gut microbiota are largely involved in host health, the present study underscored that honeybee genes may influence health directly or by developing a beneficial microbiota. Honeybees have long been regarded as a model organism for biology studies, such as social behavior, recognition, genetics and, recently, host-microbiota symbiosis[23, 88, 89]. Further identification of host alleles as shaping forces of microbial structure will advance our understanding of the host-microbe interactions.

## Conclusions

Gut microbiota affect host health and can be regulated by host genetics. We show that the gut structures are different among genetically varied bee subspecies, and the compositions dramatically shift during the social transmission among batches of founding workers and hybrid bees. Host genotype has a strong effect on the strain-level composition of *Snodgrassella*, and the bacterial Type IV pili may play an important role in microbiota-host interactions. Furthermore, we identified that the host *gluR-B* gene is associated with the abundance of *Bifidobacterium*. Mono-colonization of *Bifidobacterium* strains with a specific gene suite for polysaccharide degradation can modulate the GABA level, the expression and alternative splicing of the host genes in the brain. This work highlights the mechanisms implicated in the host-microbiota cross-talk for honeybees.

## Availability of data and materials

Sequencing data of the metagenomes has been deposited under BioProject PRJNA645015. The genomes of the bacterial isolates have been deposited under BioProject PRJNA648267. Accession numbers for genomes included in the MIDAS genomic database are provided in Dataset S1. Raw data of 16S rRNA-based amplicon sequencing has been deposited under BioProject PRJNA645267. RNA sequencing data of the honeybee brains has been deposited under BioProject PRJNA668910. The list of analysis software and scripts generated for analysis have been deposited on GitHub at: https://github.com/JiaqiangWu/bee_micro.git.

## Acknowledgments

We thank Xingan Li for help with sample collection.

## Authors’ contributions

H.Z. and H.L designed the study. J.W. generated data and performed the data analyses with contributions from X.M., Q.S., and X.H. H.L. collected samples and performed the microbiota transmission experiments. Z.Z. performed experiments with mono-colonized honeybees. H.Z., J.W., Z.Z., X.M., and X.H. prepared the manuscript.

## Supplementary Information

**Figure S1.**
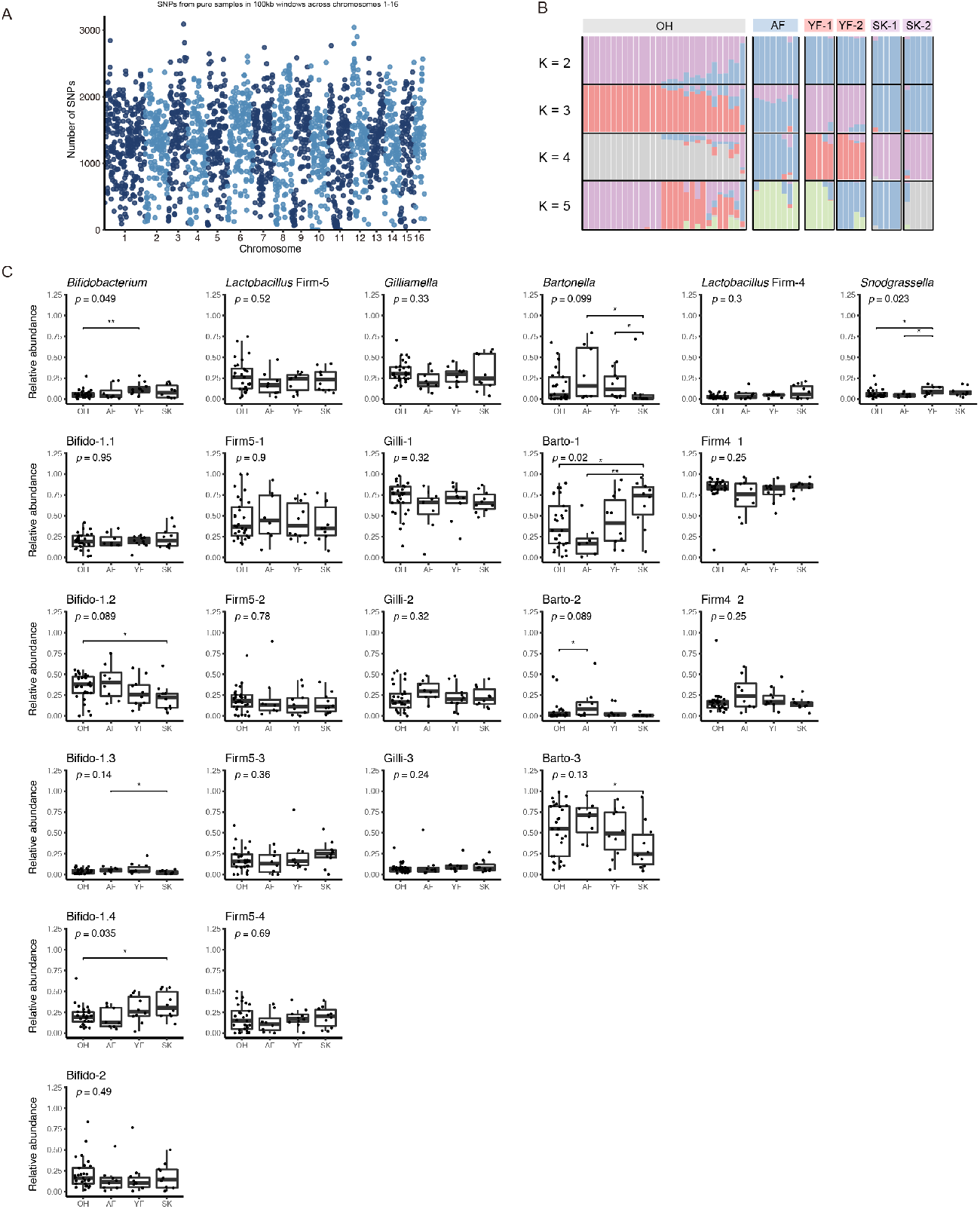
Gut community composition of genetically different purebred honeybees at both phylotype- and SDP-level. (**a**) ADMIXTURE analysis estimates the proportion of each bee genome derived from each of assumed populations (*K* = 2–5). The inferred proportion of ancestry is shown for each sample. (**b**) The number of SNPs from all purebred samples across *Apis mellifera* chromosomes 1 to 16 in 100 kb consecutive windows. (**c**) Boxplots showing the relative abundance of the core phylotypes and the SDPs from each phylotype in each subspecies group. The significance of difference among the four groups and between each pair of groups were texted (*, p < 0.05; **, p < 0.01; Wilcoxon test).

**Figure S2.**
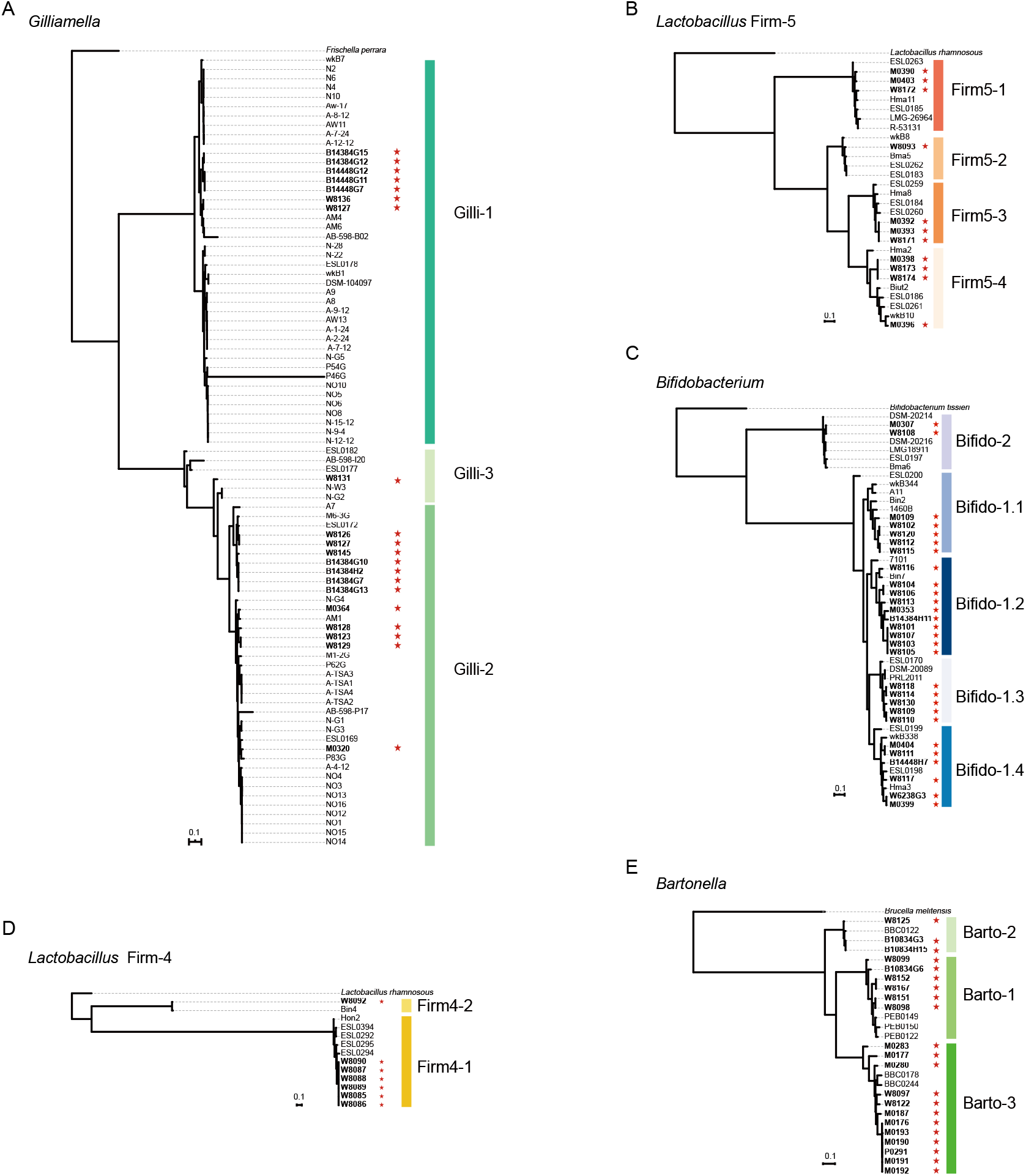
Genome phylogenies and SDP validation of the four core bacterial phylotypes and *Bartonella apis* isolated from *Apis mellifera* guts. (**a**-**f**) Phylogenies inferred for *Gilliamella* (**a**), *Lactobacillus* Firm-5 (**b**), *Bifidobacterium* (**c**), *Lactobacillus* Firm-4 (**d**), and *Bartonella* (**e**) based on the genomes of isolates in the reference database using PhyloPhlAn. Newly isolated strains in this study is annotated by a red pentagram.

**Figure S3.**
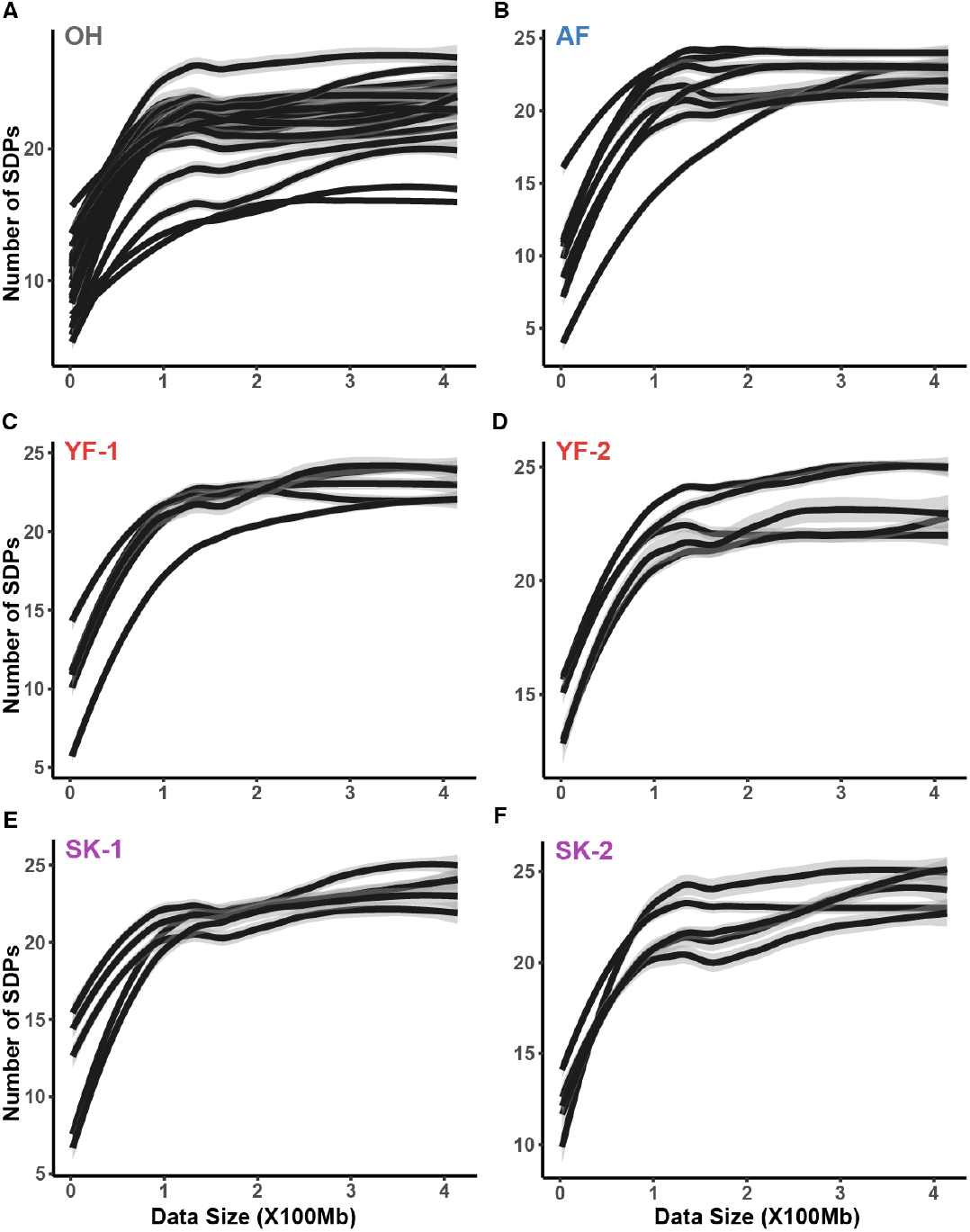
Cumulative number of observed SDPs for each metagenomic sample. The calculation of SDP richness for a given data size for each individual bee. Each panel represents one colony of a purebred subspecies, each line represents one bee individual, and the shaded area represents the 40% confidence intervals.

**Figure S4.**
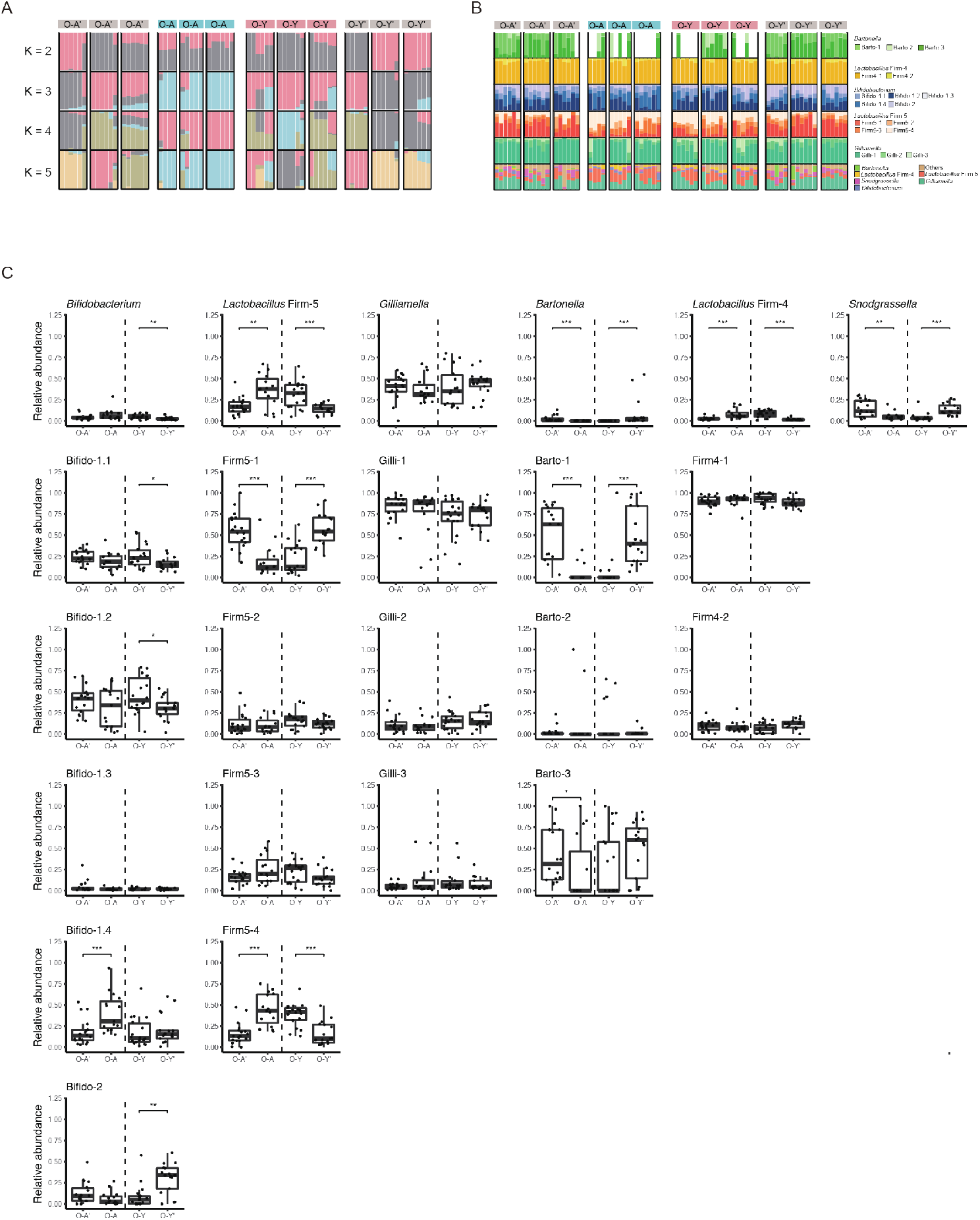
Gut community composition of founding workers and hybrid honeybees at both phylotype- and SDP-level. (**a**) ADMIXTURE analysis showing clustering of samples into 2–5 groups (*K* = 2–5). (**b**) The inferred proportion of ancestry shared with each group is shown for each sample. (**c**) Boxplots showing the relative abundance of the core phylotypes and the SDPs from each phylotype in each subspecies group. The significance of difference among the four groups and between each pair of groups were texted (*, p < 0.05; **, p < 0.01; Wilcoxon test).

**Figure S5.**
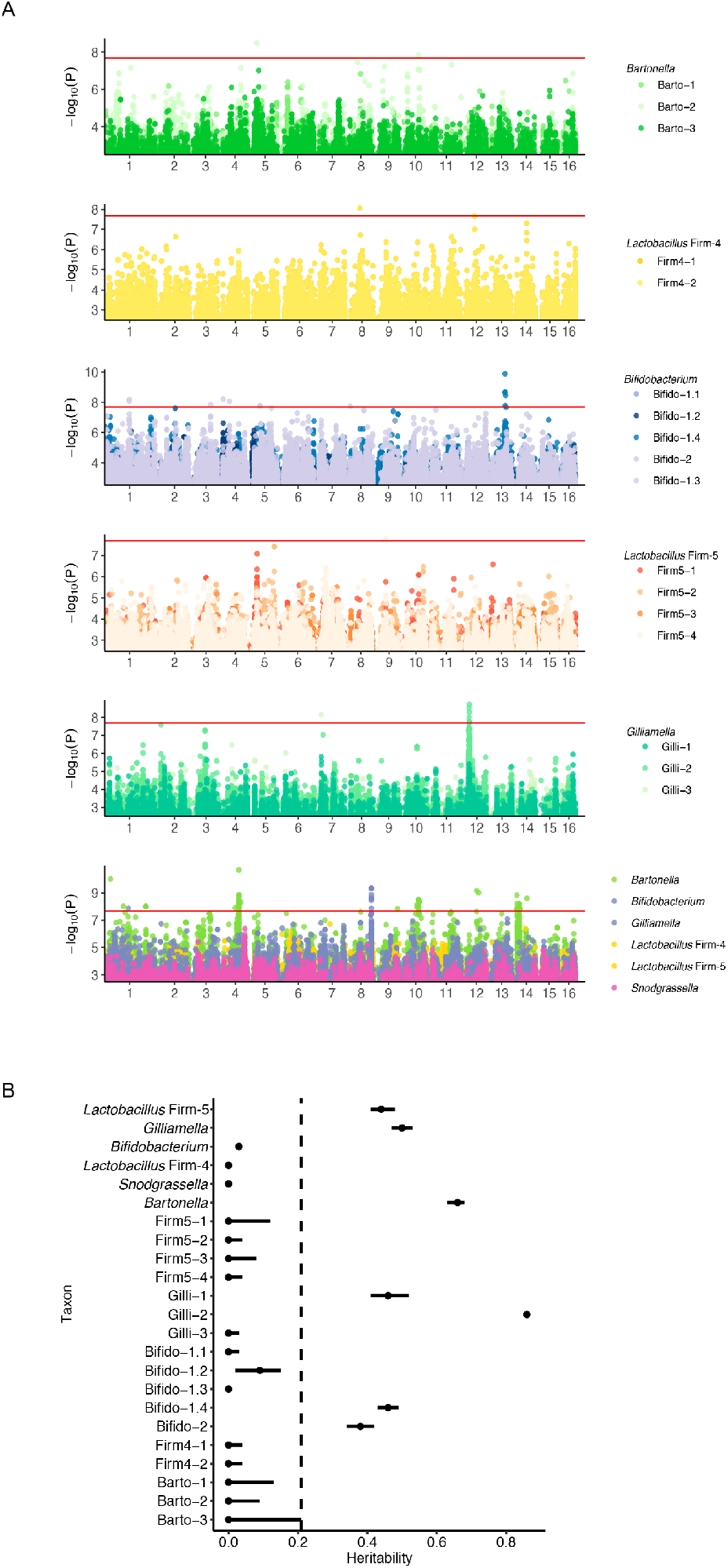
Genome-wide association analysis and heritability estimation for different phylotypes and SDPs. (**a**) Manhattan plot of genome-wide associations with different phylotypes and SDPs. The red line corresponds to a significance threshold of 2 × 10^−8^. (**b**) Heritability estimation for phylotypes and SDPs. Each point represents the heritability estimated as the proportion of variance in the relative abundances. The bars show the 95% confidence intervals around the heritability estimates.

## Additional files

**Additional file 1: Dataset S1.** The list of genomes of bacterial isolates in the database for MIDAS profiling.

**Additional file 2: Dataset S2.** Metadata of SNP distribution in Type IV pili component genes of Snodgrassella alvi in hybrid bees and founding workers.

**Additional file 3: Dataset S3.** Normalized gene expression levels in brains of microbiota-free and Bifidobacterium colonized bees.

**Additional file 4: Dataset S4.** Raw data of all metabolite abundance in the hemolymph of microbiota-free and Bifidobacterium colonized bees.

